# Distributions of LRS in varying environments

**DOI:** 10.1101/2020.06.24.169425

**Authors:** Shripad Tuljapurkar, Wenyun Zuo, Tim Coulson, Carol Horvitz, Jean-Michel Gaillard

## Abstract

Studies of lifetime reproductive success (LRS) have shown that important random events can be in ecology and evolution. Randomness should be amplified in stochastic environments, and here we show here we show this to be the case by computing the complete distribution of LRS when vital rates are Markovian can be readily computed by building on our recent paper (Tuljapurkar et al. 2020). These results complement the work of van Daalen and Caswell (2020) on moments of LRS. We use empirical studies of Roe deer, *Capreolus capreolus*, to show that environments at birth have strong effects on future performance, and that analyses of the LRS in stochastic environments are a valuable element of studies of the consequences of climate change.

## 1 Introduction

The importance of luck in determining individual lifetime reproductive success (LRS) shows how important random events can be in ecology and evolution (Tuljapurkar et al. 2009; Caswell 2011; Steiner and Tuljapurkar 2012; Snyder and Ellner 2018; van Daalen and Caswell 2017; Tuljapurkar et al. 2020). Given vital rates for a structured population (e.g., age, physiological stage, or both), moments of the LRS can be computed using a “backward” method (van Daalen and Caswell 2017), whereas a “forward” method (Tuljapurkar et al. 2020) yields the complete distribution of LRS.

A new paper by van Daalen and Caswell (2020) used the “backward” method to compute moments of LRS when vital rates vary over time but are Markovian. Complementing this analysis, we show here that the complete distribution of LRS when vital rates are Markovian can be readily computed by extending our previous work (Tuljapurkar et al. 2020). The shape of this distribution of LRS is of course strongly influenced by the probability of reproductive failure (i.e., an LRS of zero), which may result from early death or from a failure to reproduce at later ages. For the general Markov case, we also show how to compute easily the probability of reproductive failure.

We apply our new methods to empirically based estimates of age+stage vital rates for Roe dear (*Capreolus capreolus*) studied under distinct environmental conditions (Gaillard et al. 2013). We show that our method yields novel insights into the effects of an individual’s birth environment, as well as of environmental variability and serial correlation.

This paper is brief and primarily aims at making the method accessible; supporting detail is in the Appendix. We expect that the methods here will be useful in future explorations.

## 2 LRS distribution in a Markov environment

### 2.1 Preliminaries

Individuals in a structured population are followed in discrete time. Individuals are in one of *S* states, labeled with indices *i, j*, …. An individual’s “state” may be a combination of age and some other state variable such as size. All offspring start in individual state 1 (our methods also apply separately to offspring produced in other starting stages).

### 2.2 Transitions and LRS in a fixed environment

An individual transitions between states until death. Vital rates consist of transition probabilities between states: *u*(*i, j*) is the probability of a transition *i* ← *j*; these probabilities make a matrix **U**. Survival rates for states *j* = 1, 2, … are the column sums of **U**; death rates by stage are (1 − (survival rate)). Reproduction is described by probabilities Pr[*n*|*i*] that an individual in state *i* has *n* = 0, 1, … offspring.

The LRS is the random total of all offspring produced, and Tuljapurkar et al. (2020) shows how to compute the probability distribution **Γ** = {*γ*(*m*)} whose elements are the probabilities

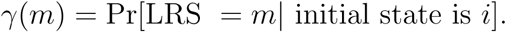

### 2.3 Variable Markovian environments

Suppose that random temporal variation in vital rates is driven by a Markovian environment. An environmental state at *t*, call it *α*, applies between *t* and *t* +1. There are *K* distinct environments (*α, β*, …) that follow a Markov chain with transition probability **P** = {*p*(*α, β*) = Pr[*α* ← *β*]}. We assume that this Markov chain is irreducible, aperiodic and has an equilibrium state in which the frequency of enviroment *α* is some *π*_*α*_ *>* 0.

In environment *α*, an individual’s stage-transition probabilities are given by **U**_*α*_ = {*U*_*α*_(*i, j*) = Pr[*i* ← *j*|environment is *α*]}.

Reproduction by an individual in stage *i* depends on the environment *α*, so there are probabilities Pr[*n*|*iα*] that this individual makes *n* offspring. Offspring are born into individual state 1, and some environmental state *α*; so an individual’s birth “state” is a combination (1 *α*).

### 2.4 Distribution of LRS: Extending Previous Results

In Tuljapurkar et al. (2020) the distribution **Γ** of the random LRS, starting with birth into individual stage *i*, is obtained by following transitions among all possible individual stages, until a random but certain death. In the present setting, a “state” *x* is a combination *iα*; and we order individual state within environment: for each environment *α*, cycle over all individual stages *i* = 1, …, *S*. Any individual is born into a state that combines the individual birth stage *i* and the birth environment *α*, and so the distribution of the LRS depends on *x* = (*iα*).

All these new states are transient because environments must change over time, but an individual must eventually die. With this expanded state space we are in essentially the same setting as in Tuljapurkar et al. (2020). Suppose that *x* = (*iα*), *y* = (*jβ*) are two of these new states. To use the results in Tuljapurkar et al. (2020), we require the transition probabilities Pr[*x* ← *y*] = Pr[*i α* ← *j β*], so we can use these transition probabilities to follow sequences of events “forward” from birth to death. It turns out that such a matrix is similar (but not identical) to the Horvitz megamatrix (Pascarella and Horvitz 1998; Tuljapurkar et al. 2003) that is used to analyze stochastic dynamics in the same expanded state space.

To keep matters simple and usable, Table 1 enumerates the steps needed to compute exactly the distribution of the LRS in a random Markovian environment. The entries in that table are operational definitions of the various probabilities, of an appropriate transition matrix, and of the computational method. Mathematical details are in the Appendix.

**Table 1:** Computing the distribution of LRS.

|  |  |
| --- | --- |
| $i, j = 1, \dots, S$ | Discrete individual states. Newborns are in the state $s = 1$. |
| $\alpha = 1, \dots, K$ | Discrete environmental states. |
| $\mathbf{P} = \Pr[\alpha \leftarrow \beta]$ | Environmental state transition probabilities |
| $\tilde{\mathbf{P}} = \mathbf{P} \otimes \mathbf{I}_S = \{p(\alpha, \beta) \mathbf{I}_S\}$ | $\mathbf{I}_S$ is $S \times S$ identity, $\tilde{\mathbf{P}}$ is a matrix of $K \times K$ blocks |
| $\mathbf{U}_\alpha, 0 \leq u_\alpha(i, j) \leq 1$ | State transition probability for $i \leftarrow j$ in environment $\alpha$ |
| $\mathbf{U} = \text{diag}(\mathbf{U}_\alpha)$ | Block diagonal formed from environment-specific transition matrices |
| $d_\alpha(j) = 1 - \sum_i u_\alpha(i, j)$ | One-period death rate for individual alive in state $j$ |
| $\mathbf{d}_\alpha$ | Column vector of above death rates for fixed environmental state $\alpha$ |
| $\mathbf{d}$ | Column vector of blocks $\mathbf{d}_\alpha$ for the $K$ environments |
| $\Pr[n\vert i\alpha]$ | Probability that an individual produces $n$ offspring when in state $i$ with the environment in state $\alpha$ |
| $\kappa_\alpha(i, z) = \sum_{n \geq 0} \Pr[n\vert i\alpha] z^n$ | Generating function for the above probabilities |
| $N$ | Integer, maximum number of offspring produced by an individual over a lifetime |
| $\theta(y) = \exp\left[\frac{-2\pi y \mathbf{i}}{N}\right], y = 0, 1, \dots, (N-1)$ | Frequencies, with $\mathbf{i} = \sqrt{-1}$ |
| $\hat{\kappa}_\alpha(i, \theta)$ | Fourier transform of $\kappa_\alpha(i, z)$: in generating function set $z = \theta$ |
| Loop over $y = 0, \dots, (N-1)$ | Given $y$, compute $\theta(y)$ |
| $\hat{\kappa}_\alpha(i, \theta(y))$ | Given $y$, compute the corresponding Fourier transform |
| $\hat{\kappa}_\alpha = \text{diag}(\hat{\kappa}_\alpha(i, \theta(y)))$ | Given $y$, a diagonal matrix for fixed $\alpha$ and all individual states, size $(S \times S)$ |
| $\tilde{\mathbf{W}} = \text{diag}(\hat{\kappa}_\alpha)$ | Given $y$, a diagonal block matrix of the preceding with $(K \times K)$ blocks |
| $\mathbf{V} = \widetilde{\mathbf{W}} \left( \mathbf{I} - \widetilde{\mathbf{P}}^T \mathbf{U}^T \widetilde{\mathbf{W}} \right)^{-1} \mathbf{d}$ | Given $y$, this computation yields a vector of length $SK \times SK$ |
| Births are in state $\{1\alpha\}$, where $\alpha$ is birth environment | Given $y$, $V(1\alpha)$ is corresponding element of vector $\mathbf{V}$ |
| For births in $\{i\alpha\}$, loop over $y$ | For each value of $y$, get a quantity $V(i\alpha)$ |
| Inverse Fourier transform the $V(i\alpha)$ | Get the $N$ values $\gamma(y)$ of the probability that the LRS equals $y$ |
| Final result is $\mathbf{\Gamma} = \{\gamma(y)\}$, $y = 0, 1, 2, \dots$ | Vector of probabilities of LRS |

The final result is the complete distribution **Γ** = *γ*(*m*) of the probabilities that an individual produces *m ≥* 0 offspring, conditional on a starting stage (1 *α*). The example in the next section shows how the entries of Table 1 are used in a realistic and complex case.

### 2.5 Final method: Probability of reproductive failure

There is a simpler way of finding *γ*(0) = Pr[LRS = 0| start in *iα*]. In an age-structured population, this probability is close to the probability of juvenile death, except e.g., for humans practicing contraception, or species with social rank that affects reproduction. In the many species where reproduction depends only on traits such as size and not age, and in general structured population models, it would be nice to compute *γ*(0) directly. We can do this by adapting the procedure above as in Table 2.

**Table 2:** Computing the distribution of LRS.

|  |  |
| --- | --- |
| Follow line 1-9 of Table 1 |  |
| $h_\alpha(i) = 1$ | For all non-reproductive individual states (e.g., juveniles, or small plants) in environment $\alpha$ |
| $h_\alpha(i) = \Pr[0\vert i\alpha]$ | Probability that an individual produces NO offspring when in potentially reproductive states in environment $\alpha$ |
| $h_\alpha = \text{diag}(h_\alpha(i))$ | A diagonal matrix for fixed $\alpha$ and all individual states, size $(S \times S)$ |
| $\widetilde{\mathbf{W}}_1 = \text{diag}(h_\alpha)$ | A diagonal block matrix of the preceding with $(K \times K)$ blocks |
| $\mathbf{v} = \widetilde{\mathbf{W}}_1 \left( \mathbf{I} - \widetilde{\mathbf{P}}^T \mathbf{U}^T \widetilde{\mathbf{W}}_1 \right)^{-1} \mathbf{d}$ | This computation yields a vector of length $SK \times SK$ |
| Births are in state $\{1\alpha\}$, where $\alpha$ is birth environment | The number $v(1\alpha)$ is corresponding element of vector $\mathbf{v}$ |
| For births in $\{1\alpha\}$ | Probability of LRS =0, $\gamma(0)$ is the number $v(1\alpha)$ |

#### Effects of changing temperatures on Roe Deer, *Capreolus capreolus*

Climate change often produces warmer temperatures and earlier springs in temperate areas, and these markedly influence the population dynamics of Roe deer, *C. capreolus*. With a fixed normal spring environment, vital rates for an age+size model based on Plard et al. (2015) were used in Tuljapurkar et al. (2020) to find the distribution of the LRS. In contrast, in early spring years Gaillard et al. (2013) found that prime-aged Roe deer females (from 1 to 7 years of age) have only 90% of the survival rates for normal spring years. Here we take “normal spring” years and “early spring” years as two environmental states, with transitions between them occurring randomly over time.

The individual states for Roe deer use 12 age classes (yearlings, 2-7 years old, 8-11 years old and *>* 11 years) and 200 size stages (200 equal body mass intervals from 1 to 44 kg). Our previous work (Tuljapurkar et al. 2020) applies only for fixed conditions in a normal spring (births into size class 100, Fig. 1, upper panel, blue solid line), or fixed conditions in an early spring (same size at birth, Fig. 1, lower panel, red solid line). Clearly a “good year” means normal spring conditions.

**Figure 1:**
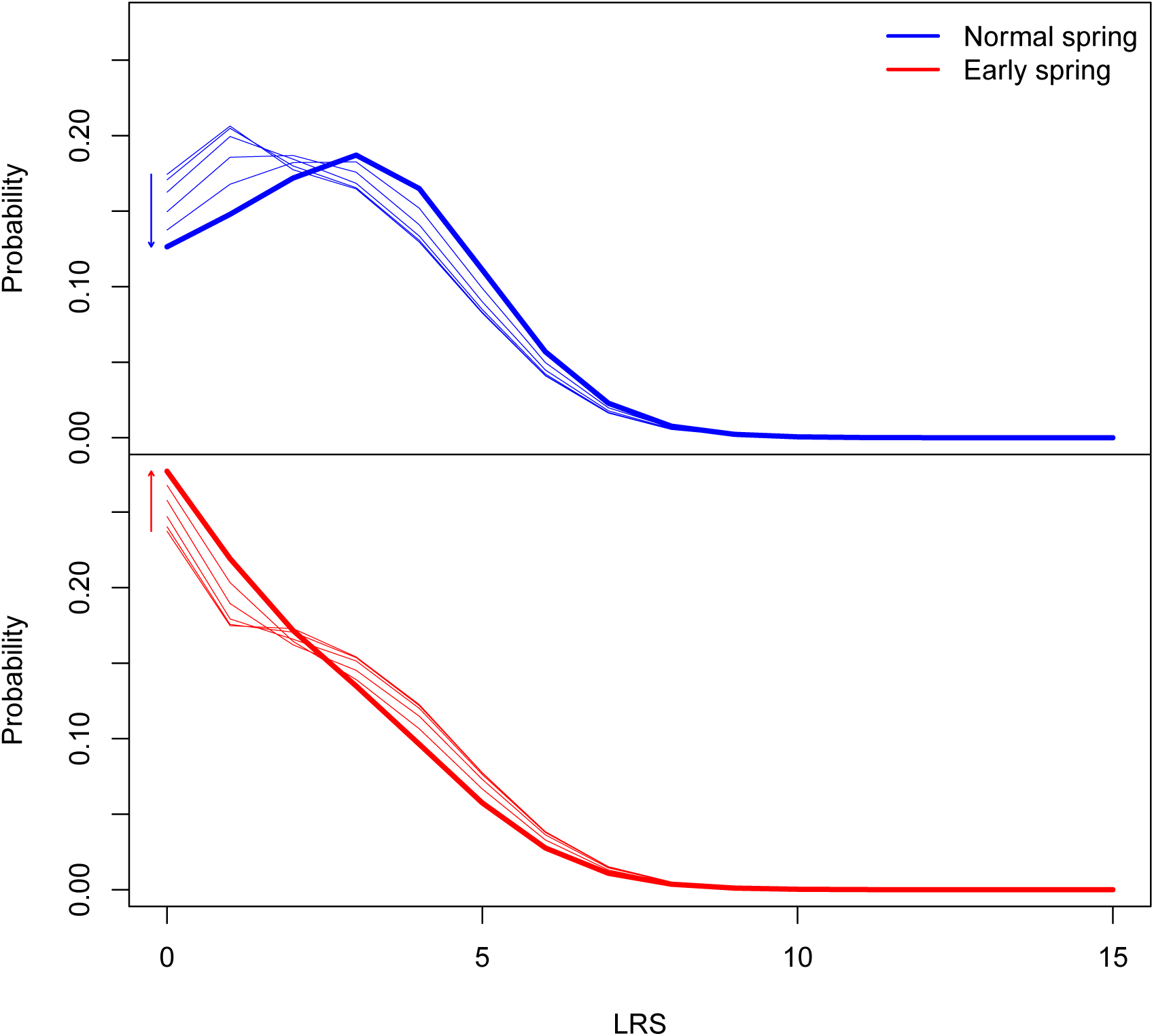
The probabilities of values of LRS for roe deer born into 2 distinct environments (normal vs. early spring) at the same initial size (stage 100). The long term environmental frequency is 50% normal spring. Autocorrelations follow the arrows with *ρ* = −0.8, −0.5, 0, 0.5, 0.8 and 1. The upper (lower) panels: probability distribution of LRS for individuals born in an early spring year (a normal spring year). In each panel, the solid line indicates a fixed environment of that type. Environmental autocorrelations follow arrows with *ρ* = −0.8, −0.5, 0, 0.5, 0.8 and 1.

We describe Markovian environmental transitions using two parameters. One is the long-run frequency *π*_1_ of environment 1 which is a normal spring; the equilibrium frequency of an early spring, environment 2, is *π*_2_ = 1 −*π*_1_. The second parameter is the serial autocorrelation, *ρ*, with value between −1 and +1, which describes the “runs” of the current environmental state. Thus, *ρ* = 0 means that the random environment is independent of the past, positive *ρ* means that the environment is “sticky” and is likelier to be last year’s environment than not, and of course, negative *ρ* means that environments are likely to alternate.

To begin, suppose that normal and early spring are equally likely (*π*_1_ = *π*_2_ = 0.5). Applying the method in Table 1, we can ask: how does LRS change with the birth environment (1 or 2), and with autocorrelation? Recall that for fixed environments, the LRS distributions (for the same size at birth) are shown by solid lines in the upper and lower panel of Fig. 1. Even with no change in the long-run frequency, the LRS distribution for either birth environment changes markedly with autocorrelation. Consider individuals born into environment 1 (upper panel): as *ρ* ranges from −0.8 (environments tend to cycle) to +0.8 (environments tend to persist), the most probable LRS for these individuals changes from near 1 to near 3. By contrast, birth into environment 2 means (lower panel of Fig. 1 that the most likely LRS is always zero regardless of environmental pattern; however the shape of the distribution changes noticeably. Clearly the birth environment matters to individual life cycles.

Suppose now that we change both *π*_1_ (the long-run frequency of a good, normal spring) and the autocorrelation *ρ*. For the same size at birth but different birth environments, our methods yield the corresponding distributions of the LRS. From these distributions we computed the average and the variance of the LRS, and the probability of reproductive failure. The averages are plotted in the upper panel of Fig. 2), with blue circles for birth environment 1, red crosses for birth environment 2. The variances are plotted in the lower panel of Fig. 2), with blue circles for birth environment 1, red crosses for birth environment 2. In each vertical segment (segments separated by gray vertical lines) the value of *π*_1_ is indicated at the top of the figure, increasing from 0.2 at the left end to 0.8 at the right end. Within every vertical segment (so for a fixed *π*_1_), birth into a normal spring results in an average LRS that increases with autocorrelation *ρ*. In contrast, birth into an early spring results in an average LRS that decreases with autocorrelation *ρ*. Thus the difference in the averages increases dramatically with autocorrelation *ρ*, regardless of the long-run frequency. Not surprisingly, increases in the probability *π*_1_ of a normal spring result in increases in the average LRS for either birth environment and a fixed autocorrelation. In contrast to this pattern of differences, the variances do not change much within every vertical segment (so for a fixed *π*_1_), chaging *ρ*). However, both variances do increase with increases in the long-run frequency *π*_1_.

**Figure 2:**
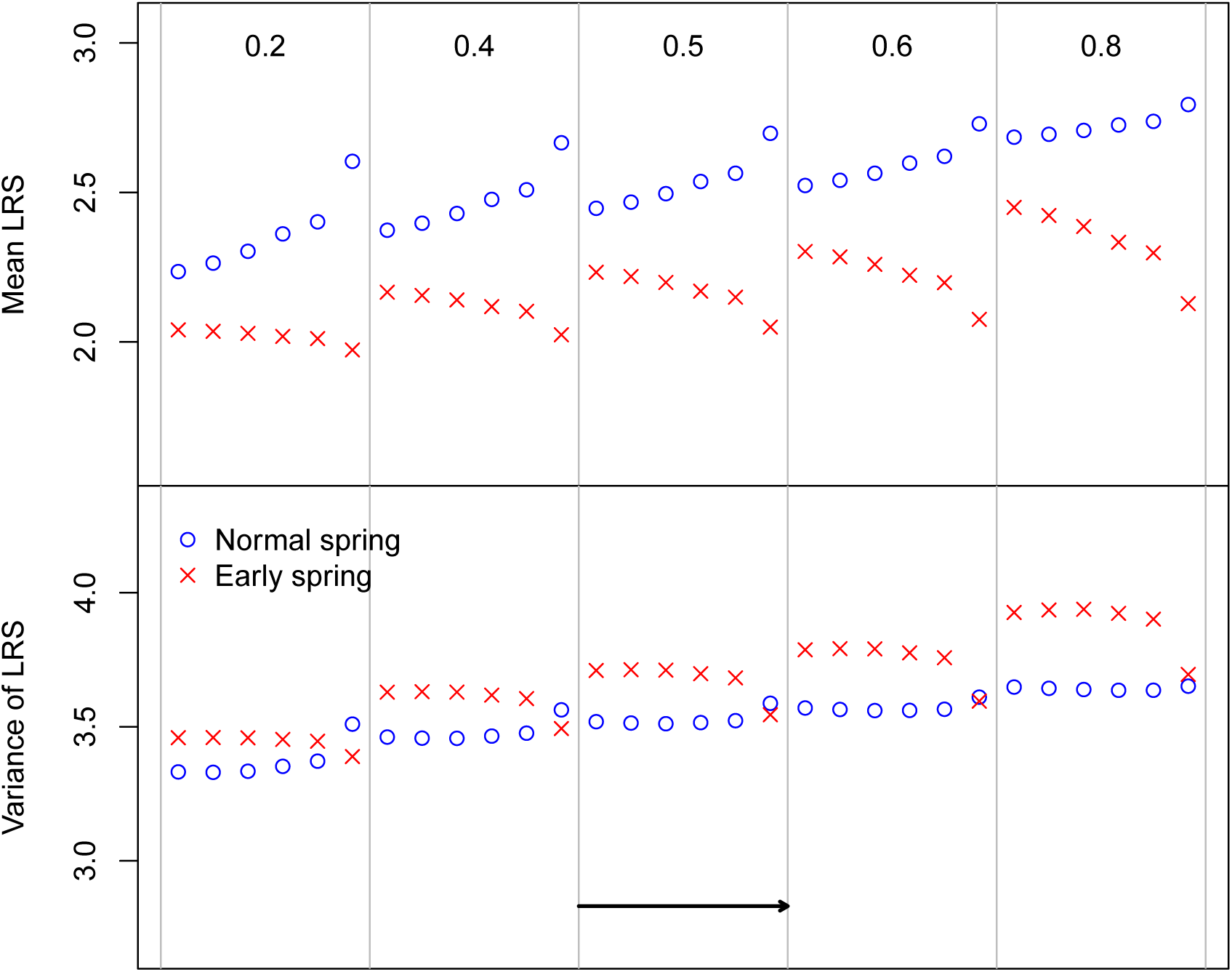
Mean (upper panel) and the variance (lower panel) of LRS for roe deer born into 2 distinct environments (normal vs. early spring) at the same initial size (stage 100). Blue circles are births in a normal spring year, red crosses are births in an early spring year. In both panels, segments shown are for the *π*_1_ marked near the top of the figure. In each segment, environmental autocorrelations follow arrow with *ρ* = −0.2, 0.0, 0.2, 0.4, 0.5, 0.8.

However, we found, but did not expect, a large difference in Pr[*LRS* = 0] for birth into a good year (environment 1) versus a poor year (environment 2) (Fig. 3). This happens because the birth year strongly affects juvenile survival for individuals born into size class 100. The difference we show here is for birth into size class 100, and decreases for much smaller birth sizes, at which juvenile survival is already so low that the added effect of birth year is modest.

**Figure 3:**
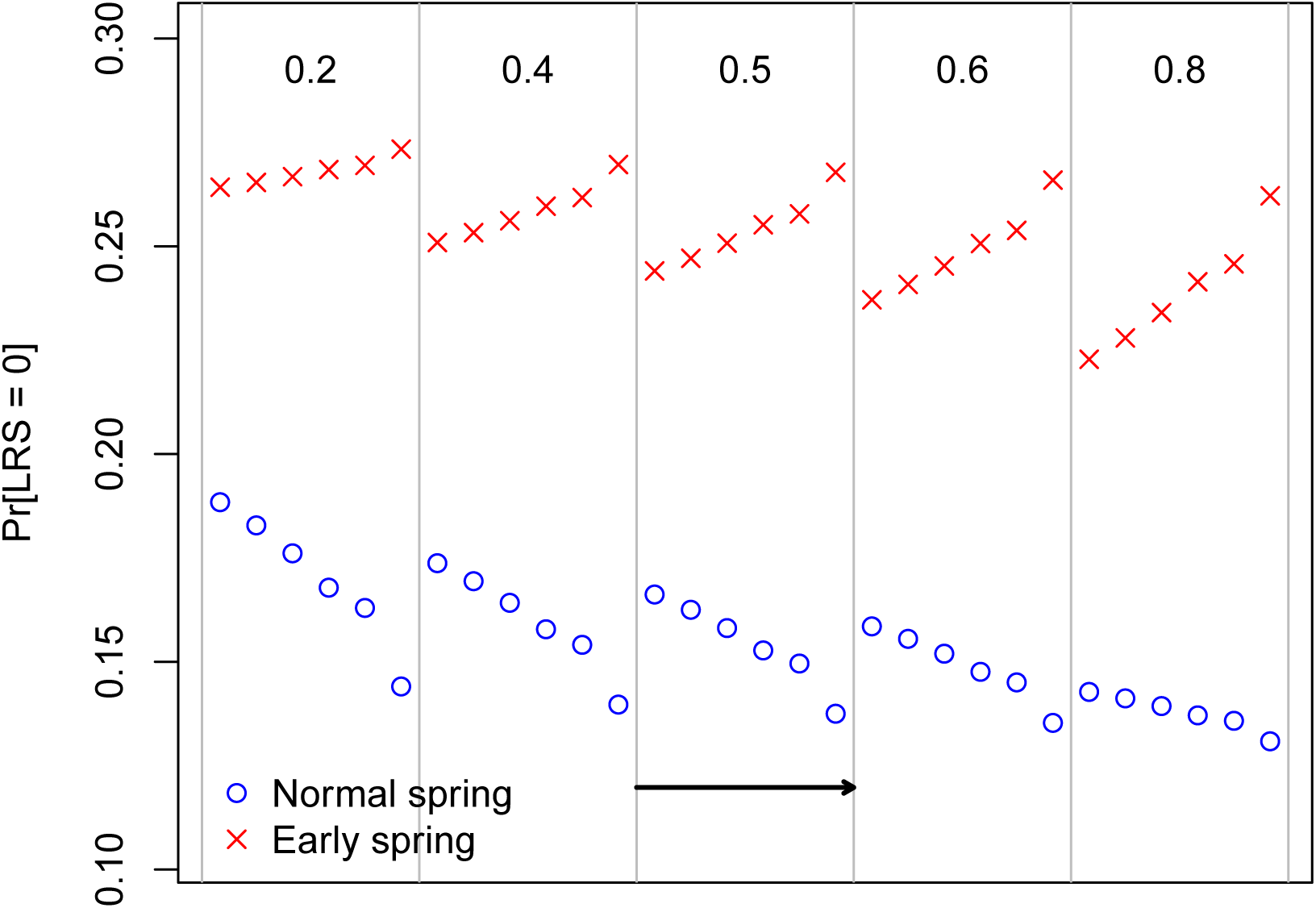
Pr[*LRS* = 0] for roe deer born into 2 distinct environments (normal vs. early spring) at the same initial size (stage 100). Blue circles, red crosses, segments, autocorrelations as in Fig. 2.

## 3 Conclusions

We have shown that the methods in Tuljapurkar et al. (2020) can readily be extended to Markovian stochastic environments. One application of the current results is to examine the effect of environmental conditions at birth on the LRS of individuals. Our example of *C. capreolus* shows that this effect can be large. Another immediate application is to examine how climate change may affect individual success. Following the work above, a model of climate change for *C. capreolus* is that normal springs become less frequency over time, meaning that the long-run frequency *π*_1_ changes from a high to a low value. The properties of LRS under different environmental patterns are readily studied using our methods. Our results complement results on the moments of LRS in Markov environments given by van Daalen and Caswell (2020).

## Appendix

### A.1 Transition Rates

To deal with the expanded state space, we define three matrices. First, stack the individual stage transition rates, the *S* × *S* matrices **U**_*α*_ into a block diagonal matrix,

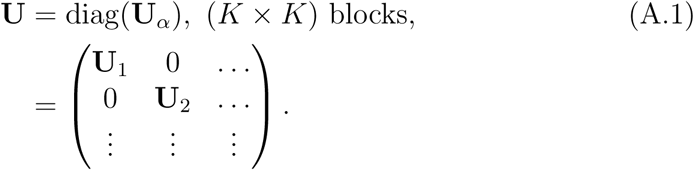

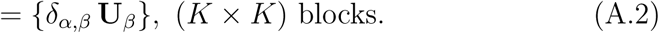

Here *δ*_*α,β*_ is Kronecker delta (see equation (A.5))

Second, use the Kronecker product × (Marcus and Minc 1992) to define a big matrix

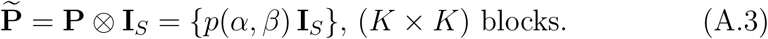

Here the matrix **I**_*S*_ is *S* × *S* identity matrix.

Third, use the preceding to make

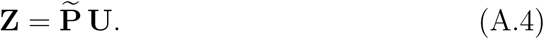

This is a matrix of *K* × *K* blocks, each of size *S* × *S*.

### A.2 Some proofs

Here and later we use the Kronecker delta:

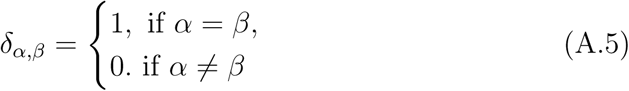

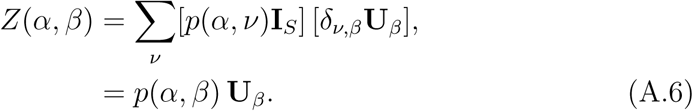

Therefore in this (*α, β*) block, the (*i, j*)th element is precisely the transition probability for both a transition (*α* ← *β*) in environmental state, and a transition (*i* ← *j*) in individual stage. Thus the joint transition probability is *Z*(*x, y*) where *x* = (*iα*), *y* = (*jβ*).

### A.3 Death Rates

The death rate for a stage *i* individual in environment *α* is just 1 minus the *i*th column sum of **U**_*α*_, **d**_*α*_(*i*) = 1 ∑_*j*_ **U**_*α*_(*i, j*).

These death rates can be made into an *S*-long vector

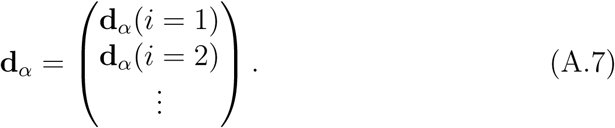

We stack these as *K* blocks into a vector that is *K S*-long, which gives us one array:

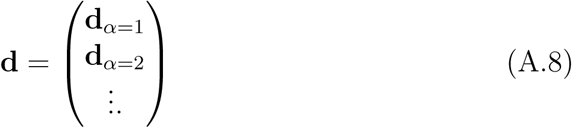

### A.4 Reproduction

This is specified by probabilities Pr[*n*|*iα*] that an individual produces *n* off-spring when in state *i* with the environment in state *α*. These probabilities are arbitrary, and may be empirical, or Poisson, or whatever. From these we have the generating functions,

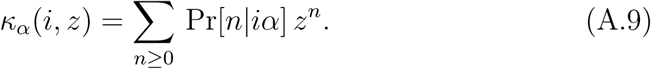

We compute the corresponding Fourier transform, 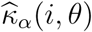, for any frequency *θ* by setting *z* = *θ*.

Fix a frequency *θ*, and collect the Fourier transforms of the pgf’s (i.e., probability generating functions) of fertility, forming a diagonal matrix of diagonal matrices:

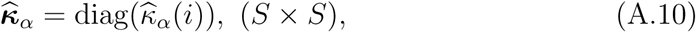

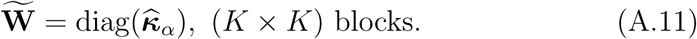

### A.5 All done

We have the matrices we need.

Using Appendix A.5 of Tuljapurkar et al. (2020), with matrix **Z** as above, we first choose integer *N*, the maximum number of offspring produced by an individual over a lifetime. Then we compute the frequencies

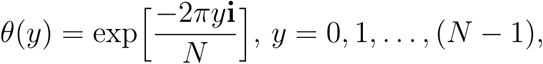

where 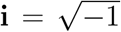. For each of these frequencies, call it *θ*, we find 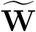 as in equation (A.11) and then compute

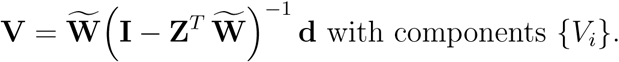

Then we can conclude that

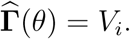

Repeat for all frequencies. Thus one computes the transform 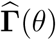.

Final step: use the Inverse FFT to find **Γ**.

